# Characterization of Wnt Signaling Genes in *Diaphorina citri*, Asian Citrus Psyllid

**DOI:** 10.1101/2020.09.21.306100

**Authors:** Chad Vosburg, Max Reynolds, Rita Noel, Teresa Shippy, Prashant S Hosmani, Mirella Flores-Gonzalez, Lukas A Mueller, Wayne B Hunter, Susan J Brown, Tom D’Elia, Surya Saha

## Abstract

The Asian citrus psyllid, *Diaphorina citri*, is an insect vector that transmits *Candidatus* Liberibacter asiaticus, the causal agent of the Huanglongbing (HLB) or citrus greening disease. This disease has devastated Florida’s citrus industry and threatens California’s industry as well as other citrus producing regions around the world. To find novel solutions to the disease, a better understanding of the vector is needed. The *D. citri* genome has been used to identify and characterize genes involved in Wnt signaling pathways. Wnt signaling is utilized for many important biological processes in metazoans, such as patterning and tissue generation. Curation based on RNA sequencing data and sequence homology confirm twenty four Wnt signaling genes within the *D. citri* genome, including homologs for beta-catenin, Frizzled receptors, and seven Wnt-ligands. Through phylogenetic analysis, we classify *D. citri* Wnt-ligands as *Wg/Wnt1, Wnt5, Wnt6, Wnt7, Wnt10, Wnt11*, and *WntA*. The *D. citri* version 3.0 genome with chromosomal length scaffolds reveals a conserved *Wnt1-Wnt6-Wnt10* gene cluster with gene configuration similar to that in *Drosophila melanogaster*. These findings provide a greater insight into the evolutionary history of *D. citri* and Wnt signaling in this important hemipteran vector. Manual annotation was essential for identifying high quality gene models. These gene models can further be used to develop molecular systems, such as CRISPR and RNAi, that target and control *D. citri* populations, to manage the spread of HLB. Manual annotation of Wnt signaling pathways was done as part of a collaborative community annotation project (https://citrusgreening.org/annotation/index).

## Introduction

*Diaphorina citri* is the insect vector of Huanglongbing (HLB, citrus greening disease), a disease that has devastated global citrus production [1,2]. HLB management is heavily based on controlling the spread of *D. citri*. In an effort to better understand the insect’s biology, the *D. citri* genome has been manually annotated to curate accurate gene model predictions. Accurate gene models can be used to develop novel insect control systems that utilize molecular therapeutics such as CRISPR and RNAi to control the spread of *D. citri* [3,4]. These molecular therapeutics would be gene-specific and reduce the reliance on broad-spectrum insecticides that have given rise to resistant *D. citri* populations [5–7].

Here, we report on *D. citri* genes involved in both canonical and noncanonical Wnt signaling. Wnt signaling is important for many biological processes in metazoans such as patterning, cell polarity, tissue generation, and stem cell maintenance [8–10]. In the model insects *Drosophila melanogaster* and *Tribolium castaneum*, knockout and knockdown of Wnt ligands and other Wnt signalling components have detrimental effects on embryo development and adult homeostasis [11–15]. Wnt signaling components could therefore serve as effective knockout targets that limit the spread of *D. citri*, reducing HLB incidence [16].

We have curated a comprehensive repertoire of Wnt signaling genes in *D. citri*. Twenty-four gene models corresponding to canonical and noncanonical Wnt signaling genes have been annotated, including seven Wnt ligands, three *frizzled* homologs, *arrow, armadillo*/*beta-catenin*, and receptor tyrosine kinases *ROR* and *doughnut*. We were unable to find *Wnt8*/*D, Wnt9*, and *Wnt16* as well as *Wnt2-4*, which have been lost in insects. The mechanisms of Wnt signaling appear to be mostly conserved and comparable to what is found in the model organism, *D*.*melanogaster* (Table 1). A model for canonical Wnt signaling in *D. citri* based on curated genes is shown (Figure 1). This is an important first step for understanding critical biological processes that may be targeted to control the spread of *D. citri* and may provide a broader insight into the mechanisms of Wnt signaling in this important hemipteran vector.

**Table 1:** Gene copy table.

| Gene | *Drosophila melanogaster* | *Apis mellifera* | *Tribolium castaneum* | *Acyrtosiphon pisum* | *Diaphorina citri* v3 |
| --- | --- | --- | --- | --- | --- |
| *Wnt1* | 1 | 1 | 1 | 1 | 1 |
| *Wnt5* | 1 | 1 | 1 | 1 | 1 |
| *Wnt6* | 1 | 1 | 1 | 0 | 1 |
| *Wnt7* | 1 | 1 | 1 | 1 | 1 |
| *Wnt8/D* | 1 | 0 | 1 | 0 | 0 |
| *Wnt9* | 1 | 0 | 1 | 0 | 0 |
| *Wnt10* | 1 | 1 | 1 | 0 | 1 |
| *Wnt11* | 0 | 1 | 1 | 1 | 1 |
| *Wnt16* | 0 | 0 | 0 | 1 | 0 |
| *WntA* | 0 | 1 | 1 | 1 | 1 |
| *pangolin* | 1 | 1 | 1 | 1 | 1 |
| *armadillo* | 1 | 1 | 2 | 2 | 1 |
| *wntless* | 1 | 1 | 1 | 1 | 1 |
| *porcupine* | 1 | 1 | 1 | 1 | 1 |
| *derailed* | 2 | 1 | 0 | 1 | 1 |
| *doughnut* | 1 | 1 | 1 | 1 | 1 |
| *arrow* | 1 | 1 | 1 | 1 | 1 |
| *frizzled* | 4 | 2 | 3 | 2 | 3 |
| *ROR* | 2 | 2 | 3 | 2 | 2 |
| *dishevelled* | 1 | 1 | 1 | 1 | 1 |
| *shaggy* | 1 | 1 | 1 | 2 | 1 |
| *Axin* | 1 | 1 | 1 | 1 | 1 |
| *ck1-gamma* | 1 | 1 | 1 | 1 | 1 |
| *Apc* | 2 | 1 | 1 | 1 | 1 |

Wnt pathway ortholog numbers in five different insect species. *Drosophila melanogaster*, *Apis mellifera*, *Tribolium castaneum*, and *Acyrtosiphon pisum* copy numbers were determined using Flybase, OrthoDB, NCBI Genbank, Uniprot, and several other publications [15,17–19]. *Diaphorina citri* numbers represent the number of manually annotated genes in the *D. citri* v3.0 genome.

**Figure 1:**
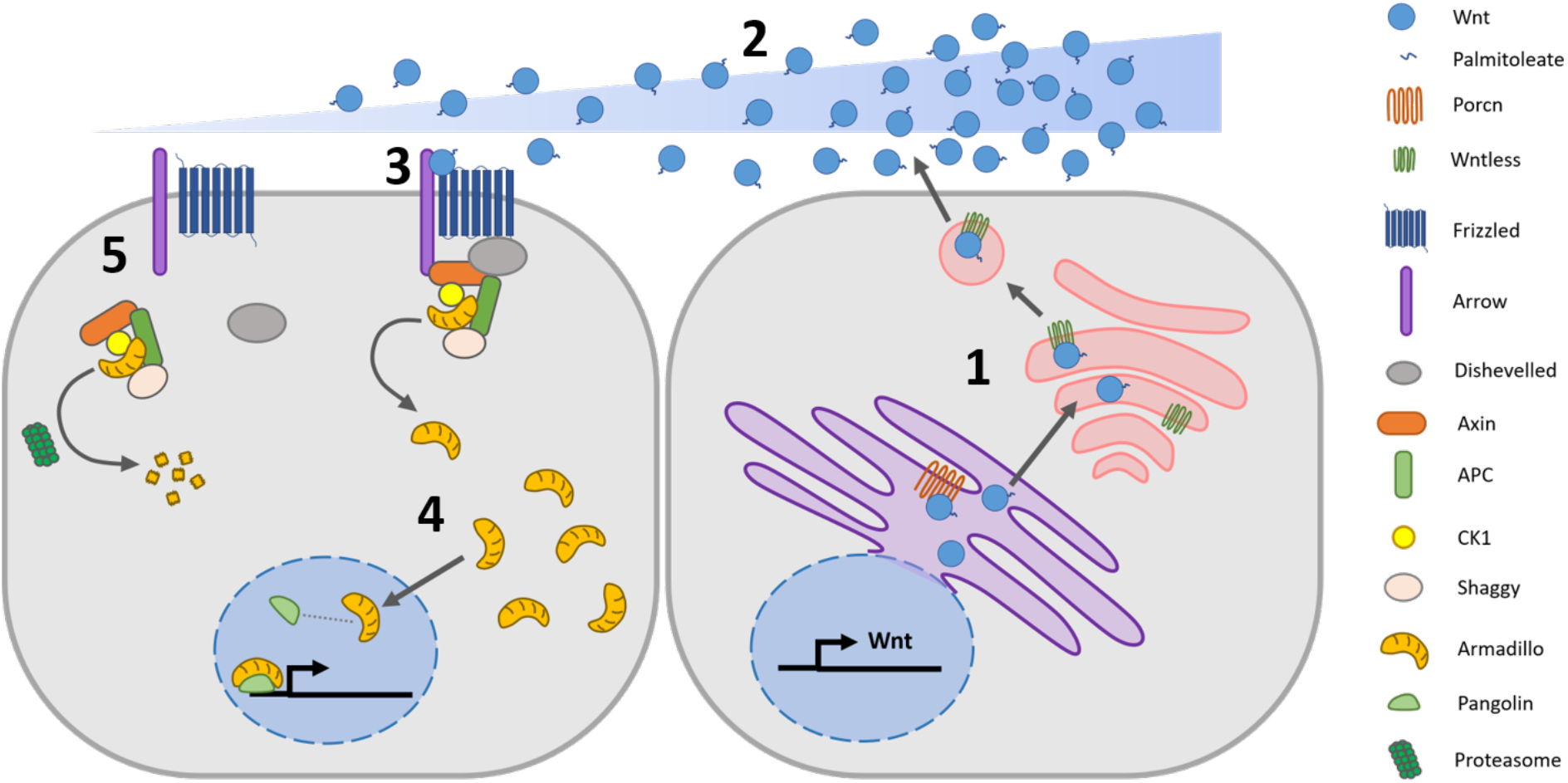
Theoretical model of canonical Wnt signaling cascade in *D. citri* based on curated genes. 1) Wnt is secreted. 2) Wnt concentration gradient forms. 3) Wnt binds to Frizzled and releases Armadillo. 4) Armadillo migrates into the nucleus, associates with transcription factor Pangolin, and regulates gene expression. 5) Armadillo is degraded in the absence of Wnt.

## Results and Discussion

The loss of Wnt ligand genes is more common in insects than in other metazoans [17], which leads to a highly variable array of *Wnt* genes and Wnt signaling components from species to species [15,18–20]. We performed a phylogenetic analysis to characterize the *D. citri* Wnt repertoire (Figure 2). Seven different *D. citri Wnts* were identified and classified as *Wnt1* (also known as *wingless*), *Wnt5, Wnt6, Wnt7, Wnt10, Wnt11*, and *WntA* (Figure 2 and 3). In comparison, seven *Wnt* genes have been identified in *D. melanogaster*, nine in *T. castaneum*, and six in *Acyrthosiphon pisum* [19,20]. The collection of *Wnt* genes found in *D. citri* is similar to other insects, and there have been no *Wnt* subfamilies identified that are unique to *D. citri*. Contrary to what has been previously reported [21], *D. citri* does appear to possess a *Wnt6* gene.

**Figure 2:**
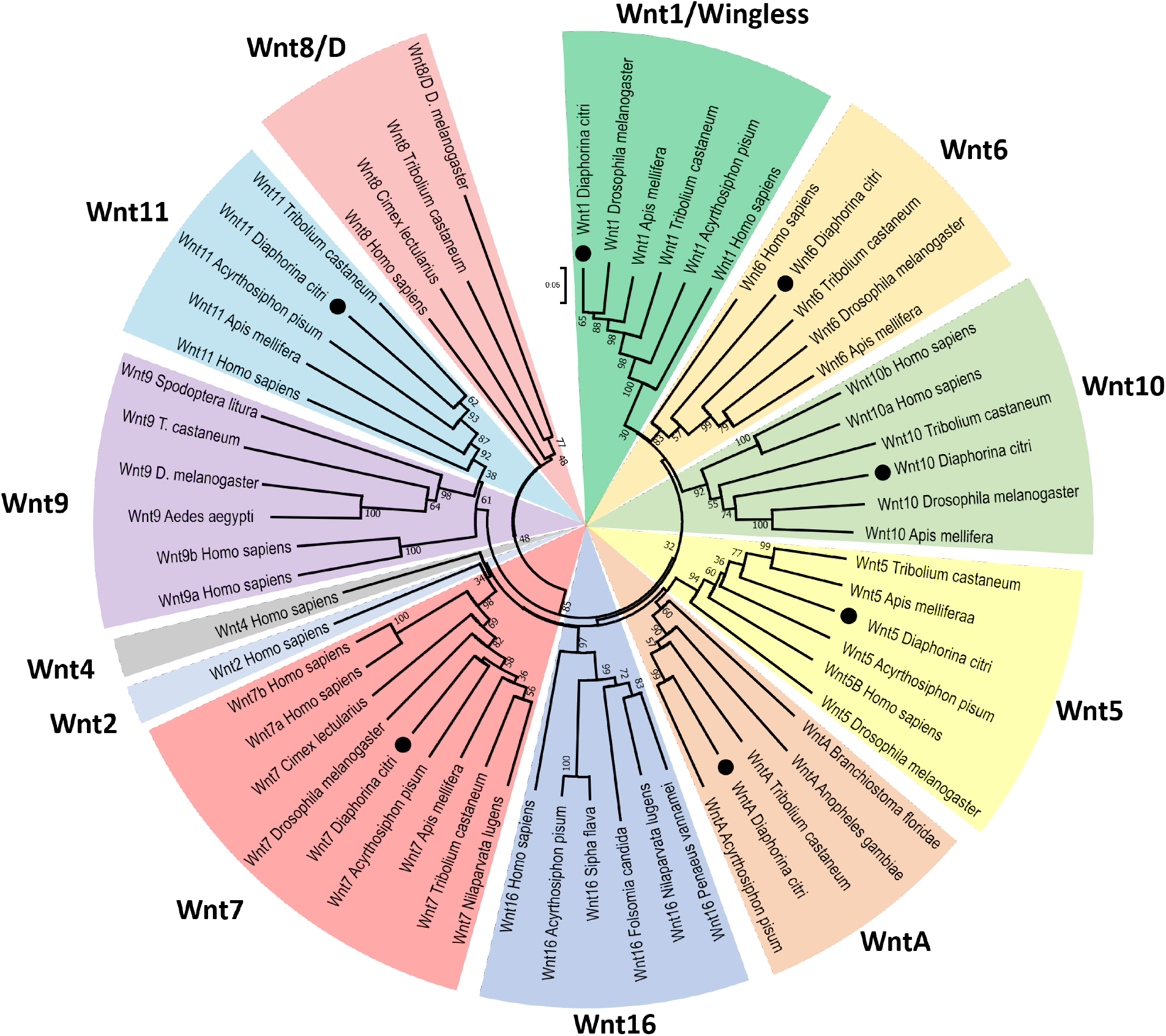
Neighbor-joining tree of Wnt protein sequences. Phylogenetic analysis was performed to categorize the seven *D. citri Wnt* genes (signified by dots). Wnt families are distinguished by clades and are color coded. Bootstrap values are based on 1000 replicates and values under 25 are removed. Ortholog sequences were collected from NCBI protein database (Table 3). Analysis was performed using MEGA7.

**Figure 3:**
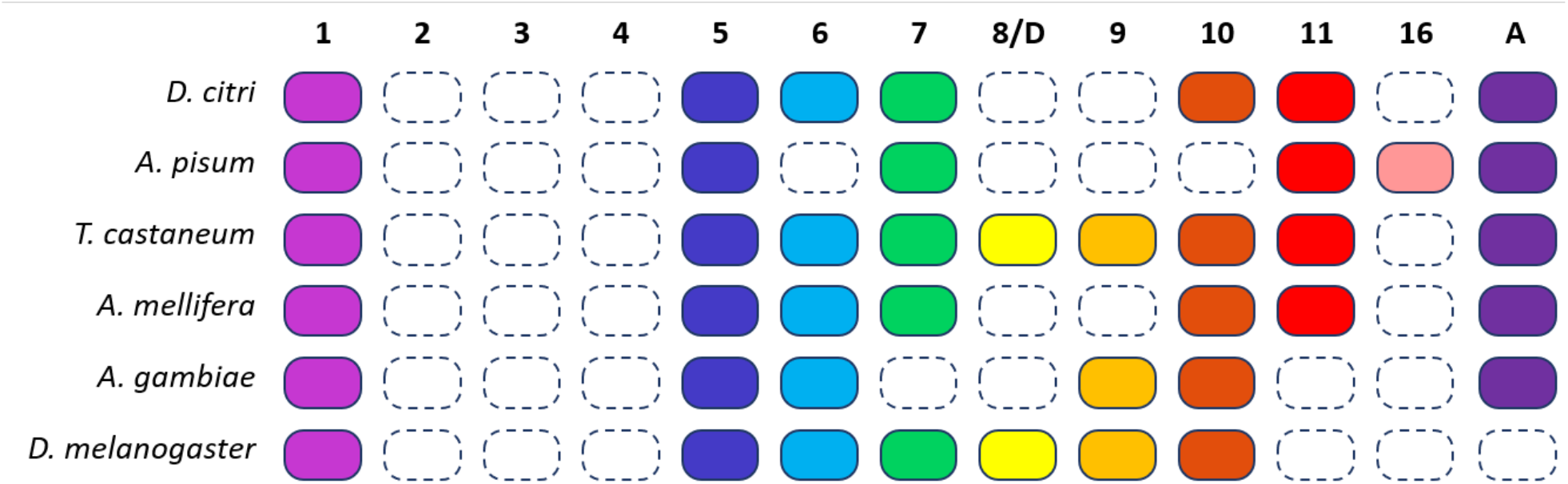
*Wnt* genes in six insects. A colored box indicates the presence of a *Wnt* subfamily (1 to 11, 16, and A) in that insect, while a white box indicates the loss of a subfamily. For example, all six species have *Wnt1* and *Wnt5*, none have *Wnt2-4*, and only *A. pisum* has *Wnt16*. Homologs of *Wnt8* in *T. castaneum* and *D. melanogaster* are also referred to as *WntD*.

*Wnt1, Wnt6*, and *Wnt10* typically occur in very close proximity in a highly conserved gene cluster [22,23]. Accordingly, it is believed that this cluster is also conserved in *D. citri* and this notion is supported by the chromosomal length genome assembly in v3.0 [24]. The close phylogenetic relationship of *Wnt1, Wnt6*, and *Wnt10* in *D. citri* (Figure 2) supports the hypothesis that this cluster is the result of an ancient duplication event, one that may predate the divergence of cnidarians and bilaterians [23]. The orientation of these clustered *D. citri Wnt* genes is similar to that found in *D. melanogaster* and differs from what may be a basal arthropodal organization of *Wnts* found in species of Coleoptera, Hymenoptera, and Cladocera (Figure 4). *Wnt9* is also associated with this gene cluster when present in the genome. However, as with *A. pisum, Wnt9* was not found in the *D. citri* genome and appears to have been lost during evolution. A second Wnt cluster, Wnt5 and Wnt7, is also common among non-insect metazoans. This cluster is not seen in *D. citri* despite the presence of both genes.

**Table 2:**
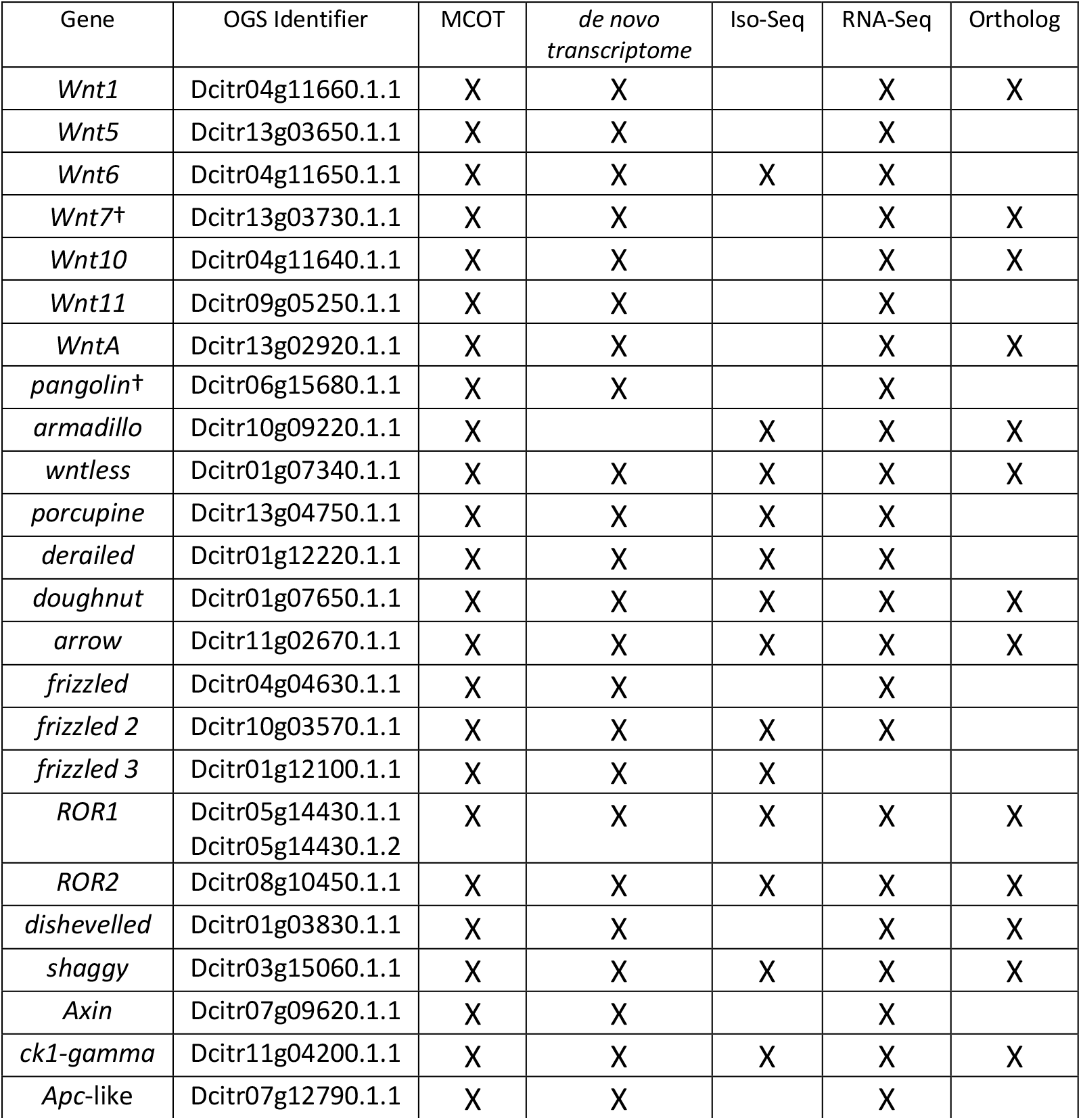
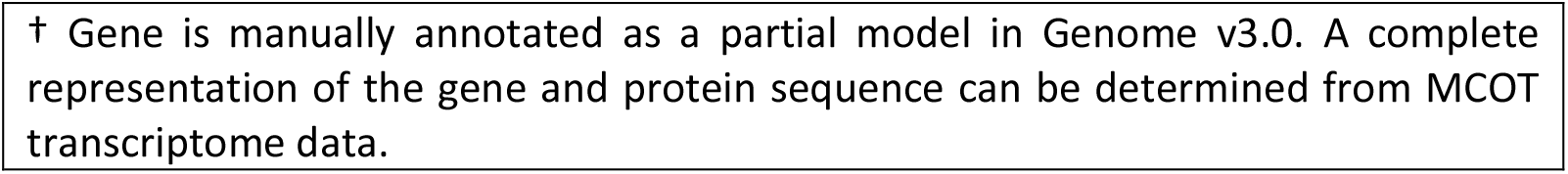
Gene Evidence Table.

| Gene | OGS Identifier | MCOT | *de novo*<br>transcriptome | Iso-Seq | RNA-Seq | Ortholog |
| --- | --- | --- | --- | --- | --- | --- |
| *Wnt1* | Dcitr04g11660.1.1 | X | X |  | X | X |
| *Wnt5* | Dcitr13g03650.1.1 | X | X |  | X |  |
| *Wnt6* | Dcitr04g11650.1.1 | X | X | X | X |  |
| *Wnt7<sup>+</sup>* | Dcitr13g03730.1.1 | X | X |  | X | X |
| *Wnt10* | Dcitr04g11640.1.1 | X | X |  | X | X |
| *Wnt11* | Dcitr09g05250.1.1 | X | X |  | X |  |
| *WntA* | Dcitr13g02920.1.1 | X | X |  | X | X |
| *pangolin<sup>+</sup>* | Dcitr06g15680.1.1 | X | X |  | X |  |
| *armadillo* | Dcitr10g09220.1.1 | X |  | X | X | X |
| *wntless* | Dcitr01g07340.1.1 | X | X | X | X | X |
| *porcupine* | Dcitr13g04750.1.1 | X | X | X | X |  |
| *derailed* | Dcitr01g12220.1.1 | X | X | X | X |  |
| *doughnut* | Dcitr01g07650.1.1 | X | X | X | X | X |
| *arrow* | Dcitr11g02670.1.1 | X | X | X | X | X |
| *frizzled* | Dcitr04g04630.1.1 | X | X |  | X |  |
| *frizzled 2* | Dcitr10g03570.1.1 | X | X | X | X |  |
| *frizzled 3* | Dcitr01g12100.1.1 | X | X | X |  |  |
| *ROR1* | Dcitr05g14430.1.1<br>Dcitr05g14430.1.2 | X | X | X | X | X |
| *ROR2* | Dcitr08g10450.1.1 | X | X | X | X | X |
| *dishevelled* | Dcitr01g03830.1.1 | X | X |  | X | X |
| *shaggy* | Dcitr03g15060.1.1 | X | X | X | X | X |
| *Axin* | Dcitr07g09620.1.1 | X | X |  | X |  |
| *ck1-gamma* | Dcitr11g04200.1.1 | X | X | X | X | X |
| *Apc-like* | Dcitr07g12790.1.1 | X | X |  | X |  |

† Gene is manually annotated as a partial model in Genome v3.0. A complete representation of the gene and protein sequence can be determined from MCOT transcriptome data.

**Table 3:** Accessions for Wnt phylogenetic tree.

| NCBI Accession: | Species: | NCBI Protein Name: | Referred to In Fig. 2 as: |
| --- | --- | --- | --- |
| XP_002609873.1 | *Branchiostoma floridae* | hypothetical protein<br>BRAFLDRAFT_60204 | WntA |
| XP_024085687.1 | *Cimex lectularius* | Wnt-8b-like | Wnt8 |
| XP_014257242.2 | *Cimex lectularius* | Wnt-7b isoform X1 | Wnt7 |
| NP_476972.2 | *Drosophila melanogaster* | Wnt oncogene analog 4<br>isoform A | Wnt9 |
| NP_476924.1 | *Drosophila melanogaster* | Wnt oncogene analog 5<br>isoform A | Wnt5 |
| NP_476810.1 | *Drosophila melanogaster* | Wnt oncogene analog 2<br>isoform A | Wnt7 |
| NP_609109.3 | *Drosophila melanogaster* | Wnt oncogene analog<br>10 | Wnt10 |
| NP_609108.3 | *Drosophila melanogaster* | Wnt oncogene analog 6<br>isoform B | Wnt6 |
| NP_523502.1 | *Drosophila melanogaster* | Wingless | Wnt1 |
| NP_650272.1 | *Drosophila melanogaster* | wnt inhibitor of dorsal | Wnt8/D |
| ALO81632.1 | *Penaeus vannamei* | Wnt-16 | Wnt16 |
| OXA45577.1 | *Folsomia candida* | Wnt-16 | Wnt16 |
| XP_025422997.1 | *Sipha flava* | Wnt-16-like | Wnt16 |
| XP_022821085.1 | *Spodoptera litura* | Wnt-4-like | Wnt9 |
| XP_015835609.1 | *Tribolium castaneum* | Wnt-4 | Wnt9 |
| XP_008196351.1 | *Tribolium castaneum* | Wnt-7b isoform X1 | Wnt7 |
| XP_008195370.1 | *Tribolium castaneum* | Wnt-1 | WntA |
| XP_015835988.1 | *Tribolium castaneum* | Wnt-11b-1 isoform X1 | Wnt11 |
| XP_008193179.1 | *Tribolium castaneum* | Wnt-10a isoform X1 | Wnt10 |
| NP_001164137.1 | *Tribolium castaneum* | Wnt6 protein precursor | Wnt6 |
| NP_001107822.1 | *Tribolium castaneum* | wingless precursor | Wnt1 |
| XP_974684.1 | *Tribolium castaneum* | Wnt-5b | Wnt5 |
| XP_971439.1 | *Tribolium castaneum* | Wnt-8a isoform X1 | Wnt8 |
| XP_021702998.1 | *Aedes aegypti* | Wnt-4 | WntA |
| XP_557821.3 | *Anopheles gambiae* | AGAP008678-PA | WntA |
| XP_006561993.1 | *Apis mellifera* | Wnt-5b isoform X1 | Wnt5 |
| XP_006557287.1 | *Apis mellifera* | Wnt-7b isoform X1 | Wnt7 |
| XP_006567803.2 | *Apis mellifera* | Wnt-11b | Wnt11 |
| XP_016771882.1 | *Apis mellifera* | Wnt-6 isoform X1 | Wnt6 |
| XP_026300091.1 | *Apis mellifera* | Wnt-1 | Wnt1 |
| XP_396944.4 | *Apis mellifera* | Wnt-10b | Wnt10 |
| XP_001949667.2 | *Acyrtosiphon pisum* | Wnt-5b | Wnt5 |
| XP_016664156.1 | *Acyrtosiphon pisum* | Wnt-16 | Wnt16 |
| XP_001948541.2 | *Acyrtosiphon pisum* | Wnt-2 | Wnt7 |
| XP_001947400.1 | *Acyrtosiphon pisum* | Wnt-1 | WntA |
| XP_001944637.3 | *Acyrtosiphon pisum* | Wnt-11b-like isoform X1 | Wnt11 |
| XP_001945295.1 | *Acyrtosiphon pisum* | Wnt-1 | Wnt1 |
| XP_022184533.1 | *Nilaparvata lugens* | Wnt-16-like | Wnt16 |
| XP_022188550.1 | *Nilaparvata lugens* | Wnt-7b | Wnt7 |
| BAB62039.1 | *Homo sapiens* | WNT5B | Wnt5B |
| NP_003382.1 | *Homo sapiens* | Wnt-2 precursor | Wnt2 |
| NP_057171.2 | *Homo sapiens* | Wnt-16 isoform 2 | Wnt16 |
| NP_004616.2 | *Homo sapiens* | Wnt-7a precursor | Wnt7a |
| NP_478679.1 | *Homo sapiens* | Wnt-7b precursor | Wnt7b |
| NP_004617.2 | *Homo sapiens* | Wnt-11 precursor | Wnt11 |
| NP_003386.1 | *Homo sapiens* | Wnt-9a precursor | Wnt9a |
| NP_003387.1 | *Homo sapiens* | Wnt-9b isoform 1 precursor | Wnt9b |
| NP_110388.2 | *Homo sapiens* | Wnt-4 precursor | Wnt4 |
| NP_079492.2 | *Homo sapiens* | Wnt-10a precursor | Wnt10a |
| NP_003385.2 | *Homo sapiens* | Wnt-10b precursor | Wnt10b |
| NP_006513.1 | *Homo sapiens* | Wnt-6 precursor | Wnt6 |
| NP_005421.1 | *Homo sapiens* | proto-oncogene Wnt-1 precursor | Wnt1 |
| NP_001287867.1 | *Homo sapiens* | Wnt-8a isoform 1 precursor | Wnt8 |

**Figure 4:**
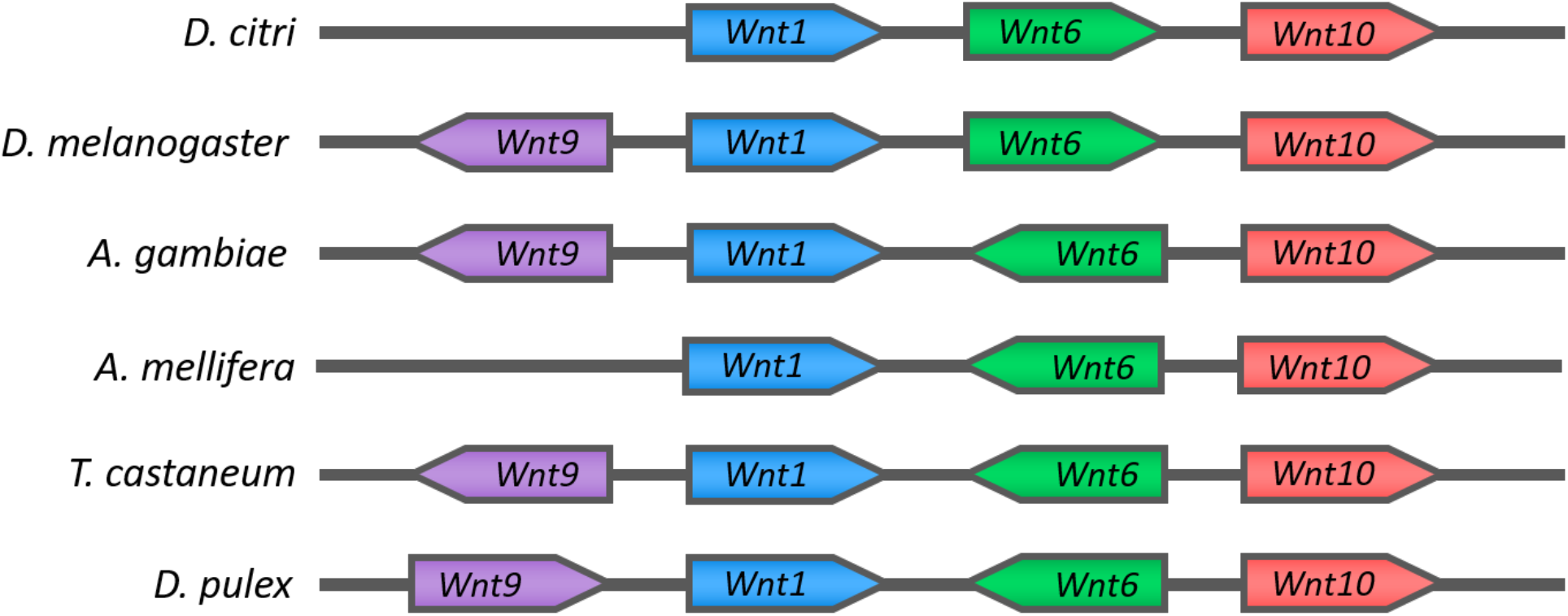
*Wnt1-6-10* Cluster comparison. Organization of *Wnt1-6-10* cluster in *D. citri* is similar to *D. melanogaster* and differs from what may be a basal arthropod gene arrangement seen in *A. gambiae, T. castaneum, A. mellifera*, and *D. pulex*. Gene lengths are not to scale.

The mechanisms that act to conserve these *Wnt* gene clusters are not well understood. In the basal metazoan, *Nematostella vectensis*, clustered *Wnt* genes do not exhibit similar expression patterns or *Hox*-like collinearity [22] and may not share regulatory elements. Data obtained from the Psyllid Expression Network (PEN) [25] available on citrugreening.org shows varying levels of expression amongst the clustered genes in different life stages of *D. citri* (Figure 5). However, it appears that *Wnt1* and *10* are similarly upregulated during embryonic psyllid development and downregulated during the adult stage, and similar transcript levels of *Wnt1* and *6* are seen in the nymphal stage. This suggests there may be shared regulation dependent upon life stage. Furthermore, ordering within the clusters is subject to rearrangement (Figure4)[20,22]. This may indicate that gene directionality is not a factor in conserving this cluster. Our annotation findings support the hypothesis that the *Wnt1-6-10* cluster is being preserved through either natural selection or an unknown mechanism, and a better understanding of the regulatory hierarchy that controls *Wnt* expression might shed light on the significance of *Wnt* gene associations in the genome.

**Figure 5:**
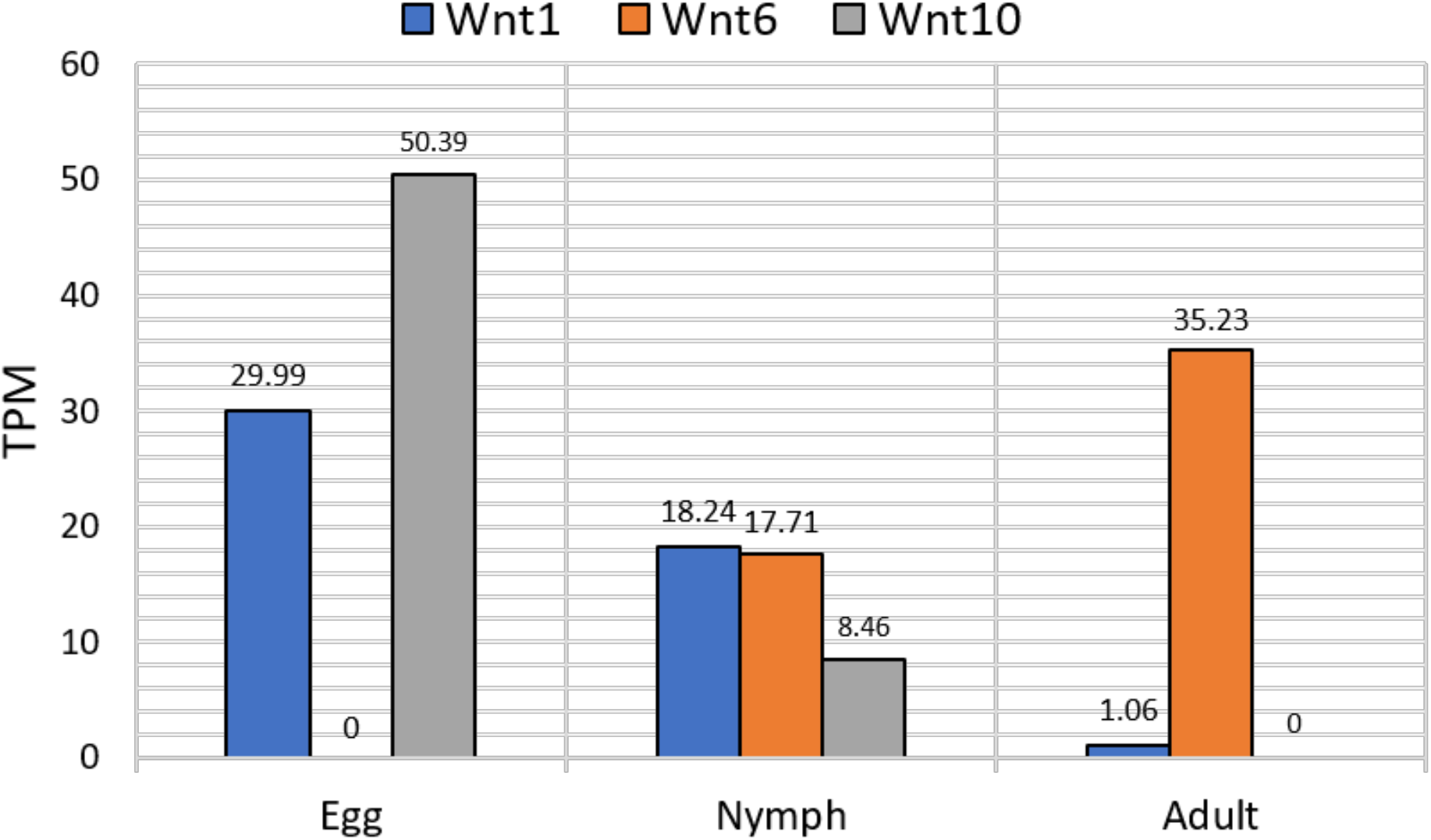
Transcript levels of clustered Wnt transcripts during different *D. citri* life stages. Whole body transcript extractions were performed on egg, nymph, and adult stages. Samples were collected from *Citrus macrophylla* and were not infected with *Candidatus* Liberibacter asiaticus. RNAseq data was collected from PEN and available on citrusgreening.org. Expression values shown in transcripts per million (TPM).

The organization of the genomic reference sequence into chromosomal length scaffolds was essential for revealing *D. citri* gene clustering. The previous genome assemblies were often unsupportive in confirming the proximity of genes due to the shorter scaffold lengths. Genome v2.0 assembly errors had likely misrepresented the location of *Wnt10*, making it appear to be separated from *Wnt1* and *Wnt6*. A complete *Wnt1-6-10* cluster was found in the improved chromosome length assembly v3.0. Thus, the quality of the reference genome should be considered when performing phylogenetic studies.

Orthologs for *Wnt2, Wnt3, Wnt4, Wnt8/D, Wnt9*, and *Wnt16* were not located in the *D. citri* genome. The close identity of certain *Wnt* subfamilies makes distinguishing between them difficult, however, the loss of *Wnt2–4* is expected as they are absent in all insects [17]. *Apis mellifera* and the hemipteran *A. pisum* have been reported to lack *Wnt8/D*, and perhaps this *Wnt* subfamily has been lost in the divergence from other insect groups [19]. Additionally, *Wnt16* was not found in *D. citri* v3.0. This finding contrasts with the gene predictions of other hemipteran genomes available at NCBI, namely *A. pisum, Sipha flava*, and *Nilaparvata lugens* (Figure 2).

Based on whole body RNA extractions collected from PEN, *Wnt6* has the highest average transcript levels of all the *Wnt* genes in both nymph and adult psyllids (Figure 6). The relatively high amount of *Wnt6* transcripts suggests that it is important during both metamorphosis and adult stage homeostasis and may serve as a good knockout target for molecular therapeutics. Transcript expression of *Wnt6* in adults is mainly concentrated in the legs and thorax, averaging 102 transcripts per million (TPM) and 272 TPM, respectively. This is considerably higher than all other *Wnt* genes in these tissues which only average between 0.26 and 3 TPM. It is unclear if other *Wnts* can be upregulated to compensate for the loss of *Wnt6*, and perhaps targeting multiple *Wnt* genes or the mechanisms by which Wnt is secreted (i.e. Porcupine and Wntless) would be more disruptive to *D. citri* physiology.

**Figure 6:**
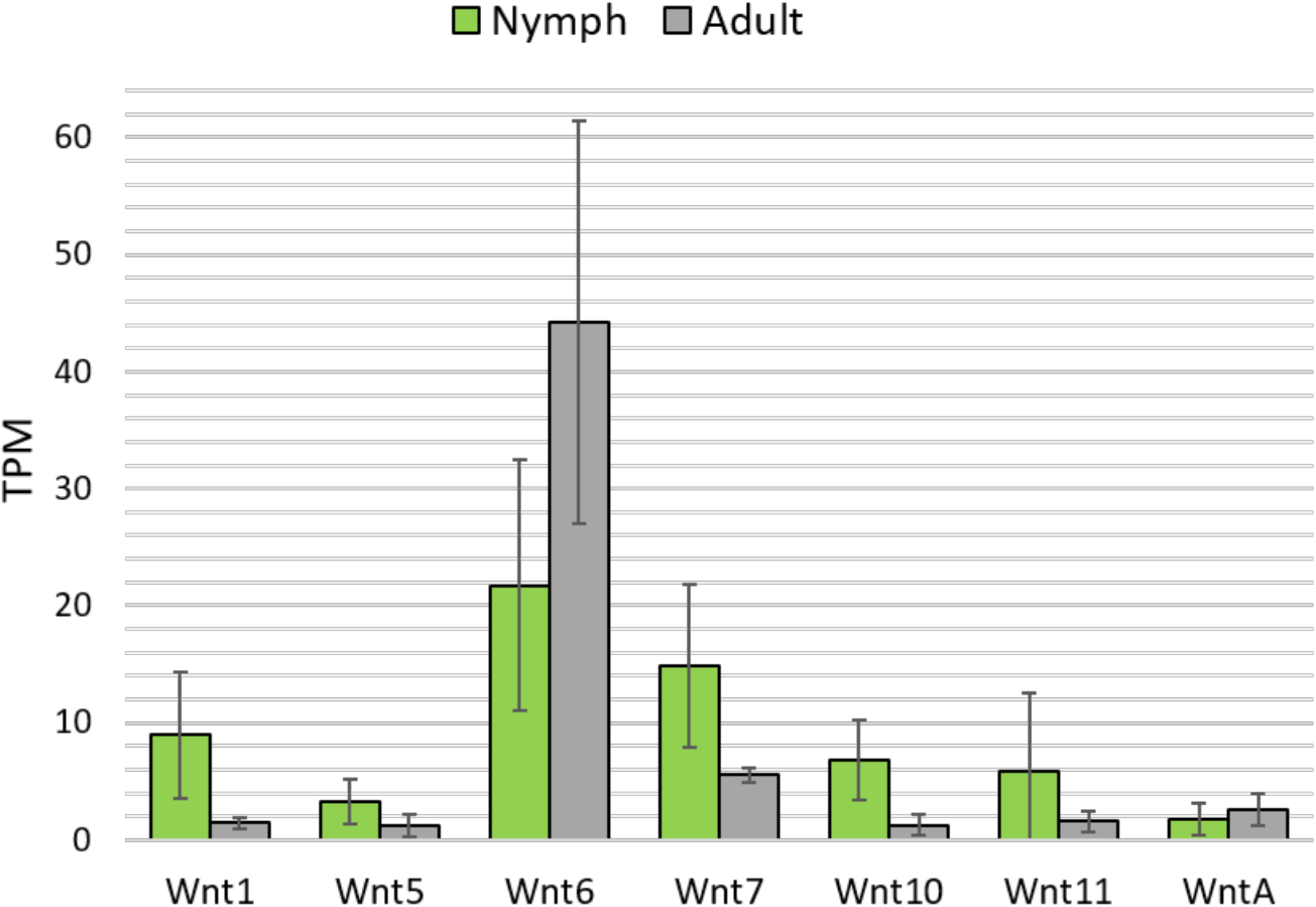
Transcript levels of *D. citri* Wnt repertoire in both nymph and adult psyllids from whole body RNA extractions. Green bars indicate the average transcript levels for *Wnt* in nymph samples, and grey bars represent the average transcript levels for *Wnt* in adult samples. Averages are based on six nymph samples and six adult samples. Expression levels shown in transcripts per million (TPM). Standard deviation of samples is shown by error bars. RNAseq data was collected from PEN and available on citrusgreening.org.

Several receptors and co-receptors associated with canonical and non-canonical signaling have been identified (Table 2). Three paralogs for the Wnt receptor encoding *frizzled* have been found in *D. citri*. We classified and numerically designated *D. citri’s* three *frizzled* genes based on how their encoded protein sequences form clades with *D. melanogaster* orthologs (Figure 7). Our analysis showed that *D. citri*, and other hemipterans such as *Halymorpha halys* and *N. lugens*, possess a Frizzled protein similar to *D. melanogaster*’s Frizzled 3. Some hemipteran Frizzled orthologs form a distinct clade separate from the Dipteran sequences (Figure 7). The hemipteran clade suggests that these genes could belong to a different subfamily of Frizzled, maybe one specific to Hemiptera, although this ortholog has not been reported in the *A. pisum* genome [19].

**Figure 7:**
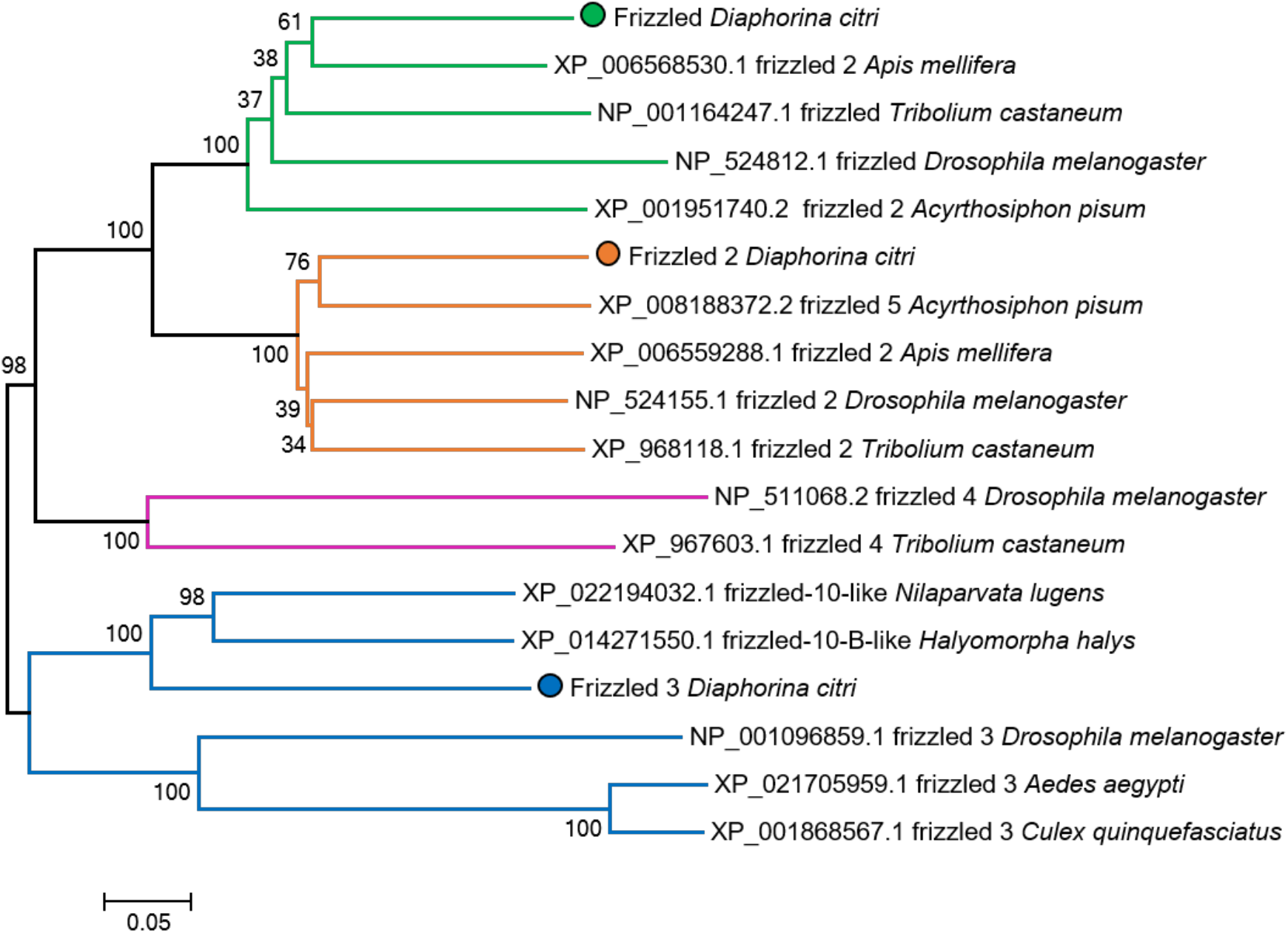
Neighbor-joining tree of insect Frizzled protein sequences. Proteins grouped in the Frizzled 1 subfamily are highlighted in green, Frizzled 2 in orange, Frizzled 3 in blue, and Frizzled 4 in magenta. Circles indicate the *D. citri* sequences. Some NCBI sequences (such as XP_006568530.1, XP_008188372.2, and XP_022194032.1) may have numeric labels derived from computational predictions that do not reflect sequence or functional similarity. Analysis performed using MEGA7.

Table of evidence supporting gene annotation. Manually annotated Wnt pathway genes in *Diaphorina citri*. Number of isoforms is noted in parentheses if there are more than one. There are 24 gene models in total. Each gene model has been assigned an identifier, and the evidence used to validate or modify the structure of the gene model has been listed. The table is marked with an ‘X’ when supporting evidence of MCOT, *de novo* transcriptome, Iso-Seq, RNA-Seq and ortholog support is present. MCOT: comprehensive transcriptome based on genome MAKER, Cufflinks, Oasis, and Trinity transcript predictions; MAKER: gene predictions; *De novo* transcriptome: an independent transcriptome using Iso-Seq long-reads and RNA-Seq data; Iso-Seq transcripts: full-length transcripts generated with Pacific Biosciences technology; RNA-Seq: reads mapped to genome are also used as supporting evidence for splice junctions; Ortholog evidence: proteins from related hemipteran species and *Drosophila melanogaster*.

Orthologs for both *ROR1* and *ROR2* have been identified. Interestingly, *ROR1* has two isoforms, the first of which contains an immunoglobulin (IG) domain that is lacking from isoform 2 (Figure 8). *ROR1* isoform 2 (Dcitr05g14430.1.2) appears to average higher transcript levels in *D. citri* egg, nymph, and adult tissues than *ROR1* isoform 1 (Dcitr05g14430.1.1) based on PEN data (Figure 9). A large number of transcripts for isoform 2 were detected in the psyllid egg (Figure 9). This suggests that expression of isoform 2 may have an important role in the early developmental stages of *D. citri*.

**Figure 8:**
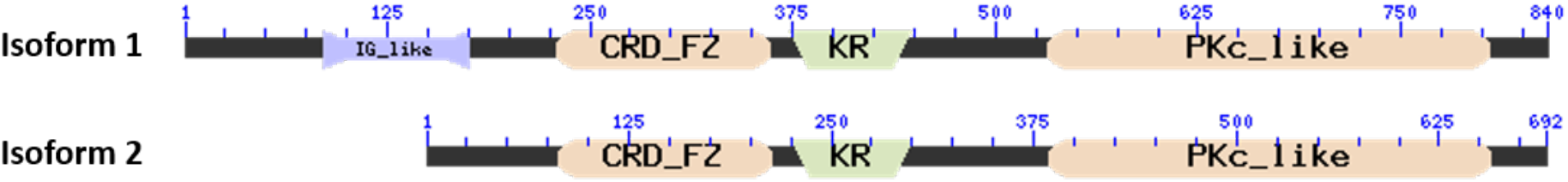
Domain comparison of *ROR1* isoforms. The immunoglobulin domain (IG_like) is present in isoform 1. Other shared domains include a cysteine-rich frizzled domain (CRD_FZ), a Kringle domain (KR), and a protein kinase catalytic domain (PKc_like). Domains were calculated and visualized using NCBI’s Conserved Domain Architecture Retrieval Tool (CDART).

**Figure 9:**
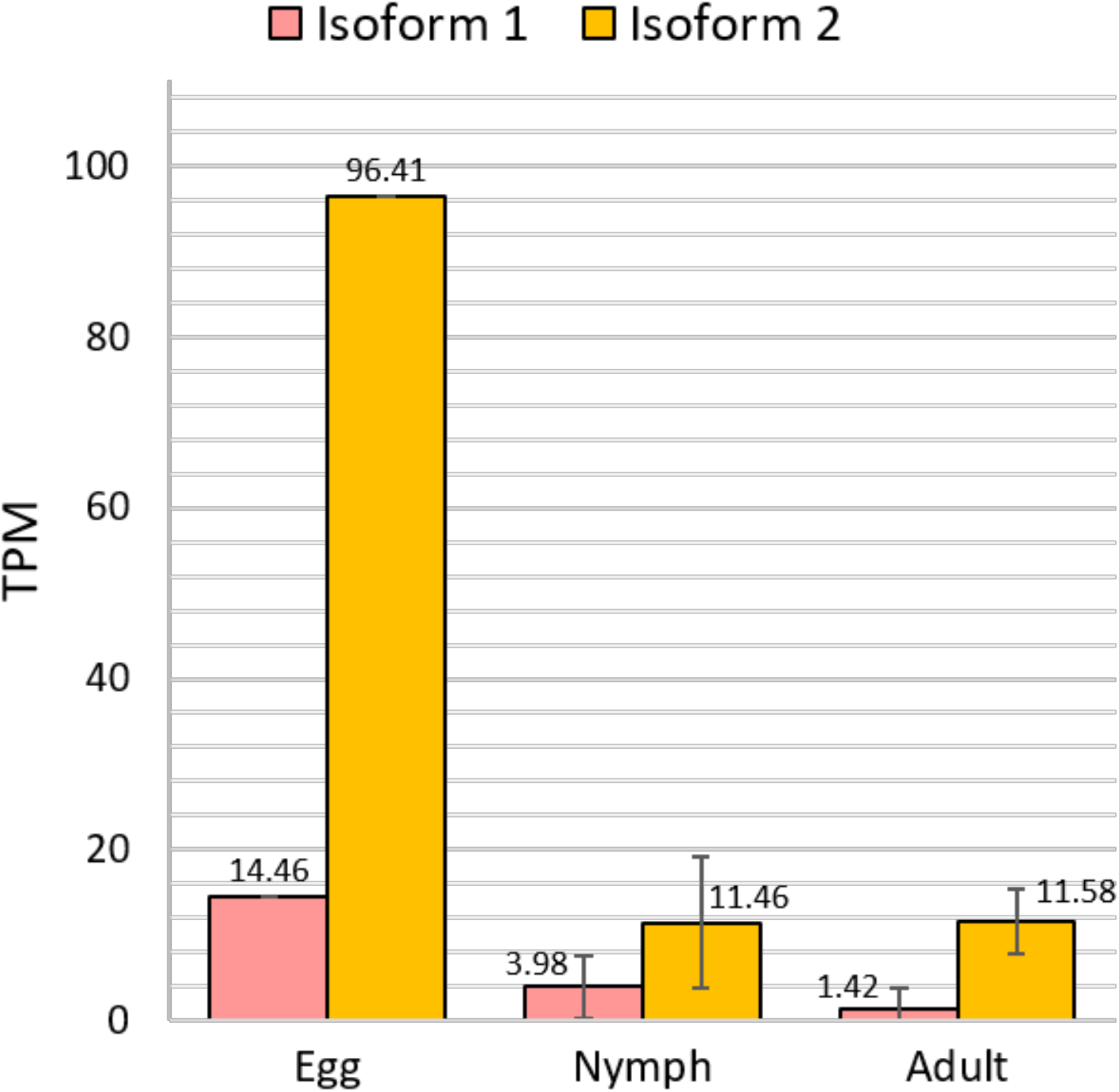
Expression of *ROR1* Isoforms in egg, nymph and adult *D. citri*. Blue bars indicate the average transcript levels for isoform 1 (Dcitr05g14430.1.1), and orange bars indicate the average transcript levels for isoform 2 (Dcitr05g14430.1.2). Note: only one egg sample was used for comparison. Egg transcripts extracted from the whole egg (1 sample total), Nymph transcripts extracted from the full body (six samples total), and adult transcripts extracted from the full body, abdomen, and thorax (14 samples total). Expression values shown in transcripts per million (TPM). Data labels note the average TPM. Standard deviation of samples is shown by error bars. RNAseq data was collected from PEN and available at citrusgreening.org.

## Conclusion

Controlling the spread of *D. citri* is an important strategy for reducing the spread of HLB. With this study we hope to provide a greater insight into *D. citri* biology as well as accurate gene models that can be used in future research and applications. We have curated a comprehensive repertoire of *Wnt* signaling genes in *D. citri*. In total, 24 gene models corresponding to canonical and noncanonical Wnt signaling have been annotated. The mechanisms of Wnt signaling appear to be mostly conserved and comparable to that which is found in *D. melanogaster* and other insects. These findings provide a greater insight into the evolutionary history of *D. citri* and Wnt signaling in this important hemipteran vector. Manual annotation and an improved genome assembly with chromosomal length scaffold were essential for identifying high quality gene models. Future work could utilize these gene models in developing CRISPR and RNAi systems that target and disrupt critical biological processes in *D. citri*, thus controlling the spread of HLB. This work was done as part of a collaborative community annotation project (https://citrusgreening.org/annotation/index).

## Methods

A complete detail of the methodology used is available at https://www.protocols.io/private/9207DE0C0FD911EBB41C0A58A9FEAC2A. To summarize, orthologous protein sequences for Wnt pathway genes were collected from the NCBI protein database and used to BLAST search the *D. citri* MCOT transcriptome database available on citrusgreening.org. The MCOT transcriptome is a transcriptome assembly utilizing Maker, Cufflinks, Oasis, and Trinity pipelines to provide a comprehensive set of predicted gene models. High scoring MCOT models were then searched on the NCBI protein database using NCBI BLAST to confirm the viability of the predicted MCOT models. The high scoring MCOT models that had promising NCBI search results were used to search the *D. citri* assembled genome. Genome regions of high sequence identity to the query sequence were investigated within JBrowse. Gene models were manually annotated using the Apollo application of JBrowse, utilizing mapped DNA-Seq, RNA-Seq, Iso-Seq, ortholog data, and other lines of evidence to edit and confirm manual annotations and gene structure. The gene models were analyzed with NCBI BLAST to assess their completeness. MUSCLE multiple sequence alignments of the *D. citri* gene model sequences and orthologous sequences were created through MEGA7 [26]. Neighbor-joining trees were constructed using MEGA7 with p-distance for determining branch length and one thousand bootstrapping replications to measure the precision of branch placement. In special cases, phylogenetic analysis in conjunction with NCBI BLAST scores was used to properly name and characterize the manually annotated gene models.

## Acknowledgements

This work was supported by USDA-NIFA grant 2015-70016-23028. We thank Alistair McGregor for valuable discussion.

